# Decomposing variation in immune response in a wild rodent population

**DOI:** 10.1101/622225

**Authors:** Klara M. Wanelik, Mike Begon, Elena Arriero, Janette E. Bradley, Ida M. Friberg, Joseph A. Jackson, Christopher H. Taylor, Steve Paterson

## Abstract

Individuals vary in their immune response and, as a result, some are more susceptible to infectious disease than others. Little is known about which components of immune pathways are responsible for this variation, but understanding these underlying processes could allow us to predict the outcome of infection for an individual, and to manage their health more effectively. In this study, we describe transcriptome-wide variation in immune response (to a standardised challenge) in a wild population of field voles (*Microtus agrestis*). We find that this variation can be categorised into three main types. We also identify markers, across these three categories, which display particularly strong individual variation in response. This work shows how a simple standardised challenge performed on a natural population can reveal complex patterns of natural variation in immune response.

## Introduction

Individuals vary in their immune response. Within a population, some individuals may fail to make protective immune responses following either natural infection or vaccination and so are especially vulnerable to infectious disease^1–4^. Defining the patterns of such variability will enhance our ability to manage the health of individuals – especially those that are most susceptible to infectious disease in human, livestock or wildlife populations.

Studies in laboratory mice are the cornerstone of immunology and have provided a detailed understanding of the molecular and cellular pathways by which immune responses are effected. This impressive mechanistic understanding, however, has only been achieved by minimising genetic and environmental variation within a laboratory setting. Where laboratory studies have examined the effects of variability – in genetics, microbiota or diet – both qualitative and quantitative differences in immune responses have been observed, with consequent effects on infection^5–7^. Nevertheless, natural variability cannot be fully reproduced in the laboratory, which has led to a recent effort to characterise the immune response in wild populations of mice or other rodents. Recent work in mice from agricultural and other anthropogenic settings is consistent with the expectation that exposure to complex environments greatly alters immune function^8^. New populations of memory T cells, present only in non-laboratory mice, have also been identified^9^.

One commonly used measure of an immune response is to assess the amount of one or more markers (e.g. transcripts or proteins) produced by a population of cells following stimulation by an immune agonist. From this *ex vivo* assay, one can gain insight into the types of immune response that could be made to a pathogen *in vivo*. Such responses depend on the cell types, the time points and the immune agonist used. Nevertheless, for any molecular marker with such a response, individuals, in natural populations especially, could exhibit different marker abundances prior to and/or following stimulation, leading to differences in their response to stimulation (here defined as the difference between marker abundances prior to and following stimulation). Furthermore, the most useful (and interesting) markers, in terms of understanding why individuals vary in their ability to mount a successful immune response, will be those for which response is most variable among individuals. In the laboratory, cell populations are usually controlled, or at least well defined, so a difference in the abundance of a particular marker can be attributed to differences in the activity of a particular cell type. However, natural variability in the abundance of a marker, and by extension in the response of individuals in the wild, could result from (i) differences in the composition of cell populations, and/or (ii) differences in the activity levels of particular cell types. Both of these components have the potential to shape the way an individual responds to immune challenge in the wild. Our intention here is not to distinguish between the two, but rather to propose a categorisation of responses, however generated.

We use a wild population of field voles (*Microtus agrestis*) to examine naturally occurring patterns of individual variation in immune response, across the transcriptome, as a first step towards furthering our understanding of the processes driving these patterns. The field population we study, in Kielder Forest Northumberland, has been the subject of extensive previous study on population ecology and pathogen dynamics^10–13^. Therefore, it allows us to place our existing understanding of more established immunological mechanisms (largely derived from the closely related laboratory mouse, *Mus musculus*) into a well-described, real-world context.

We describe three main categories of immune response: (i) uncorrelated response, (ii) constant response and (iii) baseline-dependent response (depicted in Fig. 1). We also identify markers, across these categories, which show particularly high inter-individual variability in response. We suggest that such categorisation is useful in organising natural immune variation, since little is known about which components of immune pathways are responsible for natural variability in immune response, or about the nature and possible causes of such variability. Indeed, this categorisation is not limited to the components of conventional immune pathways. The ability of an immune response to effect protection against infection, for example, will be supported by a variety of non-immune functions, that will also be activated following stimulation by an agonist, and vary to a greater or lesser extent among individuals within a natural population. By identifying the components (whether conventionally immunological or not) that are likely to be responsible for natural variability in immune response, and by describing the nature of their variability, we are laying the groundwork for exploring the processes, whether genetic or environmental, which drive inter-individual variation in immune response.

**Fig. 1.**
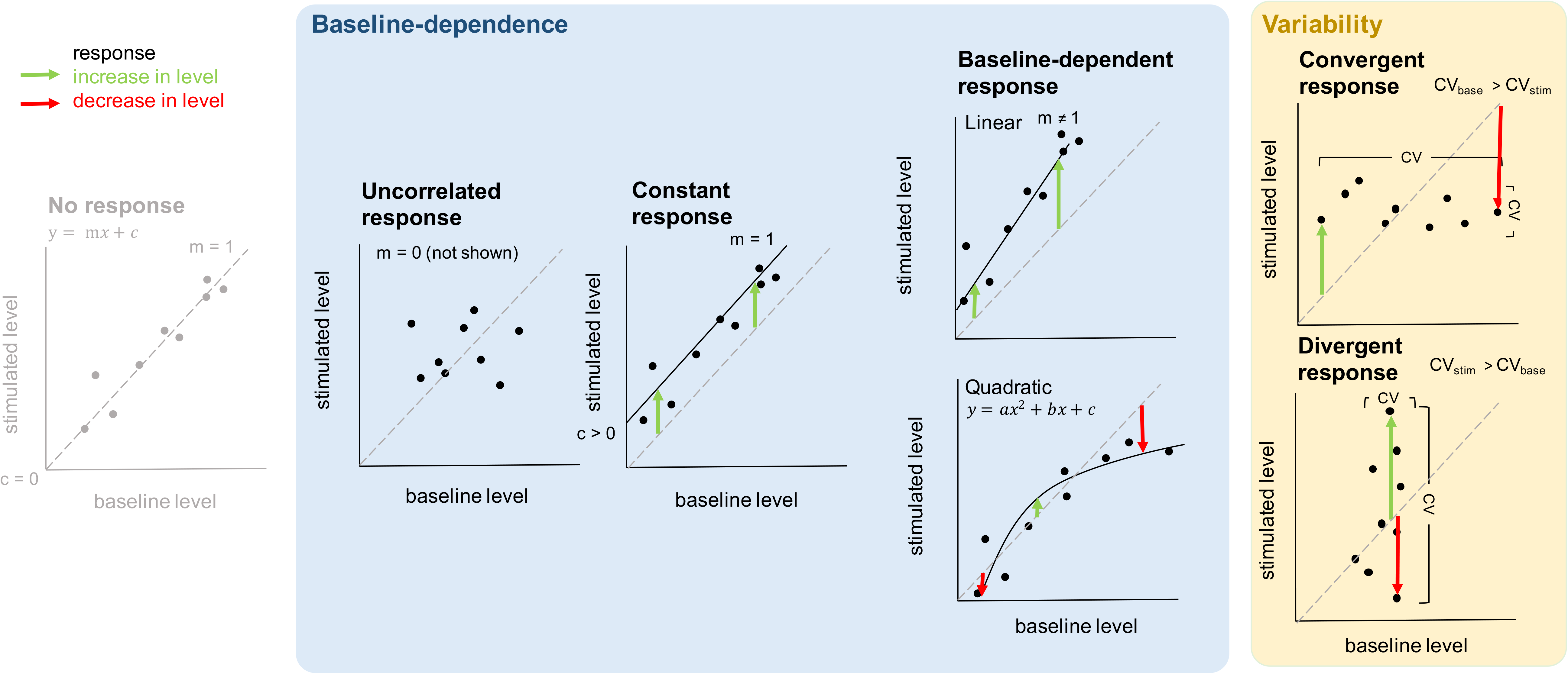
Different categories of immune response. These are based on two overlapping sets of criteria, baseline-dependence of response (blue) and inter-individual variability in response (yellow background). Arrows represent individual immune responses. No response (for reference): markers for which individuals (on average) show no response to stimulation (intercept not significantly different from zero; slope not significantly different from one). Uncorrelated response: markers for which responses of different individuals are variable and independent of their baseline level (slope not significantly different from zero). Constant response: markers for which the responses of different individuals response are (approximately) constant and independent of their baseline (intercept significantly greater than zero; slope not significantly different from one). Baseline-dependent response: markers for which responses of different individuals vary as a function of their baseline level, either as a linear function of their baseline (slope significantly different from one; slope greater than one is depicted but could equally be less than one), or as a quadratic function of their baseline (a saturating function is depicted but could equally be exponential). Convergent response: markers for which the coefficient of variation (CV) for baseline abundances is significantly greater than the CV for stimulated abundances across individuals (CV_base_ > CV_stim_). Divergent response: markers for which CV for stimulated abundances is significantly greater than CV for baseline abundances across individuals (CV_stim_ > CV_base_). Both convergent and divergent markers depicted as, but not limited to, markers for which response is uncorrelated.

## Results

### Stimulation with an immune agonist causes a widespread response

Spleen cells from sixty-two field voles were split into two populations per individual vole. One population was stimulated with anti-CD3 and anti-CD28 antibodies, while the other was kept as an unstimulated control (hereafter referred to as the baseline). 1150 transcripts (5% of all genes in the field vole genome and 85% of informative genes, those genes which were more strongly expressed; see Methods) fell into one or more of the response categories set out in Fig. 1. As expected, given that these antibodies are known to stimulate T-cell proliferation^14^, they were enriched with transcripts (hereafter markers) associated with the T-cell receptor (TCR) signalling pathway (*n* = 27; *p* < 0.001; Functional Enrichment Analysis performed in DAVID; see Methods) and other T cell-related terms: positive regulation of T-cell proliferation (*n* = 12; *p* < 0.03), TCR complex (*n* = 7; *p* < 0.001), positive thymic T-cell selection (*n* = 7; *p* < 0.01), negative thymic T-cell selection (*n* = 6; *p* = 0.03) and alpha-beta TCR complex (*n* = 5; *p* < 0.001). For the majority of these markers, a significant positive linear relationship was found between baseline and stimulated abundance (*n* = 844). Only a single marker, *Fam193b*, demonstrated a significant negative linear relationship between baseline and stimulated abundance.

### There are three main categories of immune response

Three main categories of immune response were identified based on the dependence of an individual’s response on its baseline abundance. Each of these categories demonstrates a unique pattern (Fig. 1):

#### Uncorrelated response

markers for which individuals taken from the wild differ in their baseline abundance, but the responses of different individuals are variable and independent of their baseline, such that the slope of the relationship between baseline and stimulated abundance is not significantly different from zero.

#### Constant response

markers for which individuals taken from the wild also differ in their baseline abundance, but the responses of different individuals are (approximately) constant and independent of their baseline, such that the slope of the relationship between baseline and stimulated abundance is not significantly different from one and the intercept (indicating the level of response) is significantly greater than zero.

#### Baseline-dependent response

markers for which individuals taken from the wild again differ in their baseline abundance, but the responses of different individuals vary as a function of their baseline level, either as a linear function of their baseline level (slope significantly different from one), or as a quadratic function of their baseline level, where stimulated levels either increase exponentially as a function of baseline levels or become saturated at some upper limit.

We also identified markers, across these three categories, for which variability in baseline and stimulated samples was significantly different, leading to high inter-individual variability in response (see Methods). These can be divided into two categories (Fig. 1):

#### Convergent response

markers for which variability in baseline abundance is significantly greater than variability in stimulated abundance.

#### Divergent response

markers for which variability in stimulated abundance is significantly greater than variability in baseline abundance.

### The baseline-dependent response category is most common and is significantly enriched in components of conventional immune pathways

The baseline-dependent response category was the most common (Table 1), and included a majority of markers for which stimulated levels were a linear function of baseline levels (*n* = 539), and a remainder for which they were a quadratic function (*n* = 160). The majority of quadratic response markers showed evidence for saturation (*n* = 138), indicating some upper limit on stimulated abundance. The general ontology term for immunity was enriched in the linear response category of markers (*n* = 20; *p* < 0.01). The TCR signaling pathway was enriched in the quadratic response category (*n* = 7; *p* = 0.01; Fig. 2).

**Table 1.** Table summarising the number of markers found in each of the three main categories of immune response. For each of these categories, the number of convergent and divergent markers is shown.

| Immune response category | Total no. markers | No. convergent | No. divergent |
| --- | --- | --- | --- |
| Uncorrelated | 47 | 2 | 1 |
| Constant | 306 | 0 | 91 |
| Baseline-dependent | 699 | 5 | 47 |

**Fig. 2.**
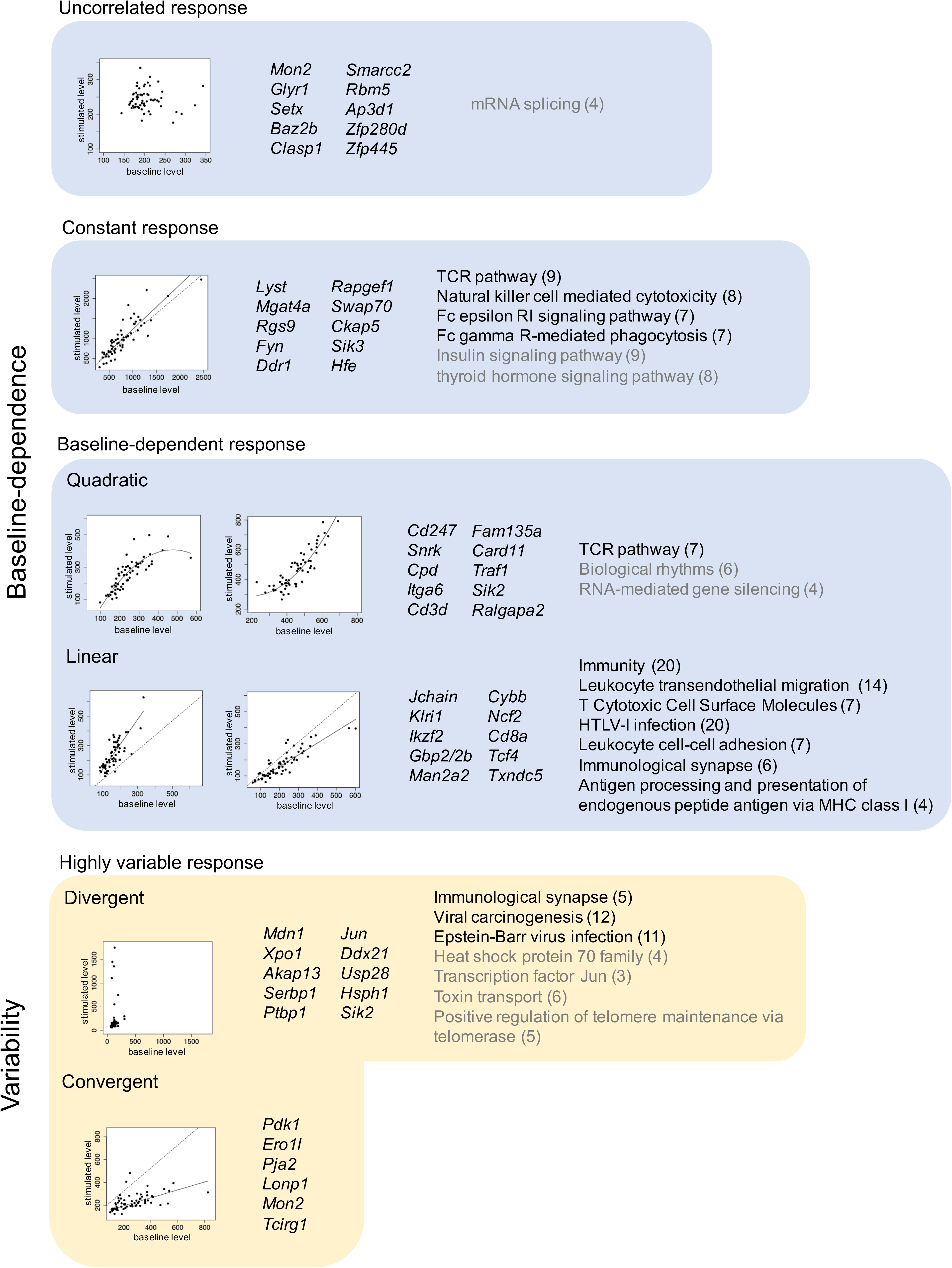
Top 10 markers and enriched ontology terms in each immune response category. Each box represents a category of immune response (as in Fig. 1). For each category, top 10 annotated markers for which we had the most confidence in their categorisation (markers were ranked on R^2^ and *p*-values) are listed, one or two of these are represented in plots showing stimulated versus baseline abundances across individuals (solid line indicates significant relationship between baseline and stimulated abundance; dashed line indicates slope equal to one for reference). In the case of the convergent category, which only included a total of six annotated markers, all markers are listed. Ontology terms of interest, from an enrichment analysis preformed on all markers within a category (where possible), are also included (immune-related terms in black).

### The uncorrelated response category is least common and lacks enrichment in components of conventional immune pathways

A number of markers showed no evidence for a relationship between baseline and stimulated abundance (*n* = 47; Table 1). For the majority of these, mean abundance was significantly greater for stimulated than for baseline samples (*n* = 39), suggesting that these markers were (on average) responding to stimulation, but to an individually variable degree, independent of baseline levels. These markers lacked any enrichment for immune-related terms (Fig. 2).

### A number of markers, including *Zap70*, show particularly high inter-individual variability in response

For a number of markers, variability in baseline and stimulated abundance was significantly different, leading to high inter-individual variability in response (*n* = 244). The vast majority of these markers showed a divergent (*n* = 237), rather than a convergent (*n* = 7) response (Table 1). Within the (stimulated) TCR signalling pathway, the highest level of variability in individual response, and the highest level of divergence, was demonstrated by *Zap70* (Fig. 3). All convergent markers fell into one of the three main immune response categories. However, over a third of divergent markers (*n* = 98), did not fall into any of these categories, appearing instead as markers which (on average) did not respond to stimulation (Table 1). Mean abundances for these markers were also not significantly different between stimulated and baseline samples.

**Fig. 3.**
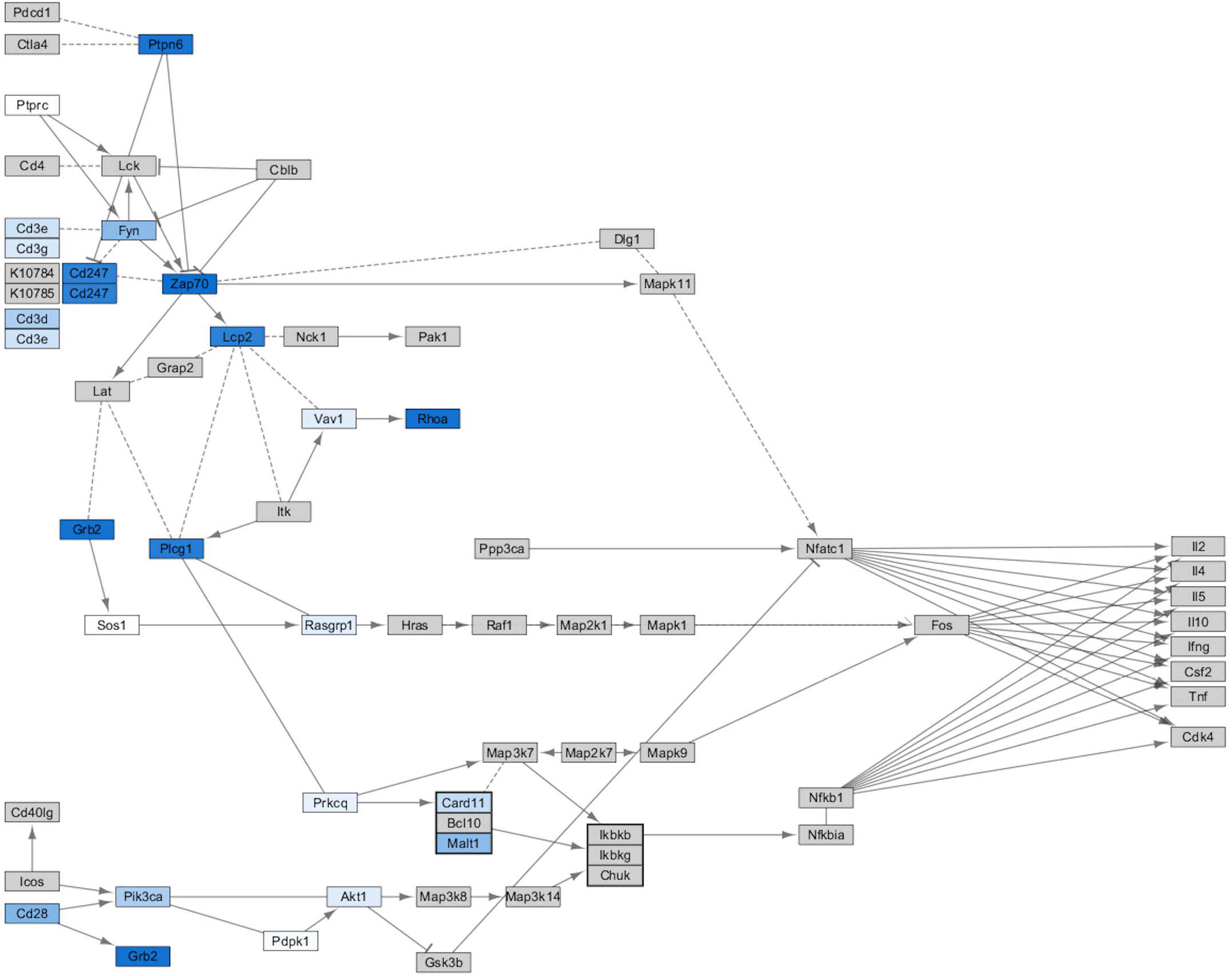
Map of the T-cell receptor signalling KEGG pathway for *Mus musculus*, with the colour of nodes representing level of inter-individual variability in response to stimulation with anti-CD3 and anti-CD28 antibodies in *Microtus agrestis*. Namely the *p*-value from an asymptotic test for the equality of variance in gene expression levels for baseline and stimulated samples (range = < 0.001–0.97). Dark blue indicates high inter-individual variability in response, whereas light blue or white indicates low inter-individual variability in response. Grey nodes represent genes for which no information is available, either because they are unannotated in the *M. agrestis* genome, or because they are weakly expressed in the spleen.

### Juveniles show more inter-individual variability in response than adults

An age-specific analysis, run separately on samples from mature (*n* = 43) and juvenile (*n* = 19) field voles, showed that higher inter-individual variability in immune response (whether divergent or convergent) was more common among juvenile voles (no. divergent markers = 108; no. convergent markers = 6) than mature voles (randomly sampled 1000 times as more samples available; mean no. divergent markers = 50, empirical 95% interval = 0–338.2; mean no. of convergent markers = 0.11, empirical 95% interval = 0–1).

### Response to stimulation is not limited to components of conventional immune pathways

Non-immune related terms were enriched in the baseline-dependent response category, including: insulin signalling pathway (*n* = 9; *p* = 0.05) and thyroid hormone signalling pathway (*n* = 8; *p* = 0.05). The top convergent response marker, *Pdk1*, is also a component of the insulin signalling pathway (Fig. 2).

## Discussion

The need to better understand variation in immune response in natural populations is now widely accepted^15–18^. Our understanding of immune responses in laboratory settings comes from animals that vary little either genetically or in prior experience. By contrast, animals in natural populations vary (perhaps extensively) in both of these. In this study, we describe natural variation in immune response in a wild population of rodents, and find that it can be categorised into a limited number of types. We identify three main categories of immune response: uncorrelated response, constant response and baseline-dependent response. We also identify markers, across these categories, which show particularly high inter-individual variability in response. Our work shows how a simple stimulatory assay performed on a natural population can reveal underlying patterns of natural variation among individuals in immune response.

The baseline-dependent response category is the largest. Markers in this category show a relationship between baseline and stimulated abundance across individuals, and their response to stimulation is (to a lesser or a greater extent) dependent on their baseline level. In some cases, individuals already expressing the greatest abundance of a marker in their natural setting went on to exhibit the greatest response to stimulation by an agonist. In others, the opposite was true, and these individuals exhibited the smallest response to stimulation. Similarly, previous work on humans has identified baseline (transcriptional) predictors of influenza vaccination response^19,20^. These differences in baseline level could be driven by either genetic variation or individual differences in past experience. In humans, genetic determinants of baseline immune cell population frequencies have been identified^21^. Even though the stimulation we describe here was not antigen specific, previous challenge by a parasite might also lead to changes in the baseline T-cell population within an individual’s spleen, affecting its response to any subsequent challenge. In fact, we find that voles infected with *Babesia microti* (a blood parasite, common in our population^22^) have larger spleens than uninfected voles^13^. This prior experience may prime an individual, enabling a greater response to subsequent challenge (e.g. slope greater than one; Fig. 1). However, individuals may also have an upper limit on the number of immue cells they have available^23,24^. An individual that is already mounting an immune response to a parasite, and has a large number of activated T cells, could therefore respond less to a similar challenge than an ‘immunologically naïve’ individual (slope less than one; Fig. 1). Membership of the baseline-dependent response category recapitulates the known biology of the immune response (being highly enriched for immune ontogeny terms). In doing so, it validates the approach we use here, as a way of identifying markers of immune significance.

In some cases, individuals varied in their natural abundance of a marker but their response was unrelated to this. They did nevertheless respond to stimulation, with the majority of these markers occurring at a significantly higher mean abundance in stimulated samples than in baseline samples. This uncorrelated response category, which contains a moderate number of markers, also lacks any enrichment for immune-related ontology terms. This suggests that markers in this category are not conventional immune markers but could be of immune significance. We warn against omitting such markers from studies of immune response in the laboratory. They could play an important part in our understanding of the immune response, indicating for example, genetic variation in response among individuals, which is independent of baseline level.

Cutting across this categorisation, a large number of markers displayed a pattern in which variation between individuals was particularly strong. We describe two types of such markers, both of which could be used in future studies as indicators of natural variability in immune response. Markers in the less common, convergent, response category showed much greater variation naturally than following stimulation. This pattern may be characteristic of markers showing variable levels of prior activation, coupled with some maximum or optimum abundance, and resulting in a stabilisation of the immune response across the population following stimulation. We found that convergent patterns were more common among juvenile voles. This could suggest that they are more constrained in the energy they have available (as a result of the competing energetic demands of growth and development) or the number of immune cells they have available (as a result of a developing immune system). Either resource constraint could result in a maximum abundance, making them more inclined to converge. Due to the costly nature of the immune response, individuals often trade-off their investment in different arms of the immune system^25,26^. Different types of immune response are therefore likely to be associated with different optimum abundances (or regions) and an individual already mounting an immune response, but to a different type of challenge (associated with different cell types), may respond by down-regulating expression.

Divergent markers, which were more common, showed much greater variation following stimulation than there was naturally. This pattern may be characteristic of (but not limited to) markers showing genetic variation in response to the agonist, independent of baseline levels e.g. subsets of animals that appear similar but respond more strongly to stimulation than others. Our own recent work, where we found an association between polymorphism in a single gene and a marker of a more tolerant immune response^27^, is an example of such genetic variation in immune response. Further supporting this hypothesis, here, we found more divergent markers among juvenile voles than mature voles. Younger voles are expected to have less variable exposure histories, as a result of their shorter life spans, making it easier to detect genetic effects. Equally, though, divergent patterns could be the result of differences in early life experiences. One would also expect these to be more easily detectable in juveniles.

The divergent category (predominantly) included markers for which individuals made (on average) the same response to stimulation and markers that did not respond (on average) to stimulation. Standard differential expression analysis would miss the individual variation present in the former group, and would fail to pick up the latter group of markers altogether. Both warn against looking at average (population-level) response, and point instead, to the value of looking at individual-level differences in immune response. This is particularly important because divergent markers may act as critical regulators of pathways. For example, *Zap70*, which demonstrates particularly high levels of variability in individual response and is centrally located in the TCR signalling pathway, interacts with many other markers (Fig. 3). We suggest that *Zap70* expression could be used as a marker of response in larger studies. Indeed, it is already linked to major seasonal immune variation in wild fish^28^ and is being used as a prognostic marker for B-cell chronic lymphocytic leukemia in humans, with potential implications for determining a patient’s treatment path (recently reviewed in Liu *et al.*^29^). Other potential prognostic (or diagnostic) factors which may have been missed using standard differential expression analyses may be present in this category and warrant further investigation.

The immune response categories we describe here are based on spleen cells stimulated with anti-CD28 and anti-CD3 antibodies and sampled at 24 hours. However, the relative frequency of the response categories reported here may vary depending on the choice of agonist and/or time point. For example, markers are known to follow different response trajectories, with some immediately responding and reaching peak activation, and others taking longer to reach this point^30^. Sampling at a later time point, then, when the ‘slower’ markers have reached peak activation, may lead to more convergence than reported here. In order to fully account for this temporal variation, multiple time points need to be averaged across. We argue that both time-specific and averaged responses are of functional significance, but hope others will extend our work. We use RNASeq here in order to give a broad view of the immune response. Single-cell RNASeq could be used to quantify differences in individual response resulting solely from differences in cell-specific activity. Previous work has shown that transcript levels generally correlate with protein levels across genes^31^. However, more work is needed to confirm these response categories at the functional level^32^. In future, Q-PCR or protein-level data could be used in order to include weakly expressed markers, which were excluded here as a result of the heteroscedasticity inherent in RNASeq data.

Markers that responded to stimulation were not limited to immune pathways as conventionally defined. They included, for example, markers involved in the insulin signalling pathway. This is in line with previous studies, which suggest that insulin plays a key role in coordinating an organism’s response to infection, influencing, in particular, the allocation of resources^33,34^. One of these markers, *Pdk1*, was also among the top convergent markers. This could be representative of the high levels of variability in the (baseline) nutritional status of individuals in a natural population, coupled with an upper limit on the processes involved in glucose metabolism.

The immune categories we presented here, therefore, highlight markers not traditionally associated with immune functions, and offer a promising avenue for identifying potential prognostic (or diagnostic) factors for disease, like *Zap70*. They also point to both genetics and prior experience as drivers of natural variation in immune response. Our future work will further decompose this natural variation into that driven by these two components.

## Methods

### Field methods

Sixty-two field voles were collected between July and October 2015 to assay expression by RNASeq. These voles came from four sites in Kielder Forest, Northumberland (55°130N, 2°330W). Each site contained a trapping grid of regularly spaced traps (at approx. 5 m intervals) and was also used for other components of a larger field study (for more details see Wanelik *et al.*^27^).

### Ethics statement

All animal procedures, carried out as part of this field study, were performed with approval from the University of Liverpool Welfare Committee and under the authority of the UK Home Office (Animals (Scientific Procedures) Act 1986) project licence number PPL 70/8210 to SP. Voles were killed by a rising concentration of CO_2_ followed by exsanguination.

### Cell culture methods

Splenocyte cultures from each vole were split into two populations, one of which was stimulated with anti-CD3 antibodies (Hamster Anti-Mouse CD3e, Clone 500A2 from BD Pharmingen) and anti-CD28 antibodies (Hamster Anti-Mouse CD28, Clone 37.51 from BD Tombo Biosciences) at concentrations of 2 μg/ml and of 1 μg/ml respectively for 24hr, and the other was left as an unstimulated control to act as a reference level. We refer to this reference level as the baseline, and control samples as baseline samples. However, it is important to note that this level will vary for an individual, not only on a day to day basis, but throughout its life. Culture conditions were otherwise equivalent to those used in Jackson *et al.* (2011)^35^. Costimulation with anti-CD3 and anti-CD28 antibodies was used to selectively promote the proliferation of T cells^14^, our assumption being that this would reflect the potential response of T-cell populations *in vivo*. Cell populations within splenocyte cultures were variable but left undefined.

### RNASeq preparation and mapping

RNA was extracted using Invitrogen PureLink kits. Following extraction, cDNA libraries were prepared using Illumina RiboZero kits and libraries were constructed with NEBNext Ultra directional RNA library prep kit according to the manufacturers protocols. Samples were sequenced to produce 2 × 75 bp paired-end reads on an Illumina HiSeq4000 platform. Adaptor sequences were removed with CUTADAPT version 1.2. and further trimmed with SICKLE version 1.200 (minimum window quality score of 20). This resulted in a mean library size of 18 million (range = 5–50 million) paired-end reads.

High-quality reads were mapped against a draft genome for *M. agrestis* (GenBank Accession no. LIQJ00000000) using TOPHAT version 2.1.0, and a set of predicted gene models was generated using BRAKER. Mapped reads were counted using FEATURECOUNTS. Further analysis was performed on counts of mapped reads for each gene in R version 3.4.2^36^. These count data were initially filtered to remove unexpressed genes (those genes with fewer than three counts per million across all samples; *n* = 13). Following filtering, library sizes were recalculated and data were normalised to represent counts per million (cpm). These data were found to be correlated with quantitative PCR (Q-PCR) data (see Supplementary Fig. 1). No correction for gene length was necessary as all analyses were based on comparisons across (rather than within) samples. Transcript abundance for a particular gene here represents a single, functional measure of its activity across some, undefined, cell population. In order to maximise the power of our analysis to identify biologically relevant patterns, we focussed on those genes which were expressed at an informative level in the spleen prior to and/or following stimulation (*n* = 1350 or 6%). Genes expressed at a mean level greater than 200 cpm were considered informative. As weakly expressed genes were removed (minimising heteroscedasticity), log-transformation of data was unnecessary (Supplementary Fig. 2).

### Statistical analysis

Genes for which a response to stimulation was observed across individuals were identified, and, as elaborated in the Results, categorised on the basis of (i) the dependence of an individual’s response on its baseline level, and (ii) the degree of inter-individual variability in response across individuals.

#### Baseline-dependence of response

The dependence of an individual’s response on its baseline level was quantified by testing the relationship between that individual’s baseline abundance (cpm_base_) and its stimulated abundance (cpm_stim_) using a linear regression, taking the form

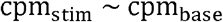

as well as a quadratic regression, taking the form

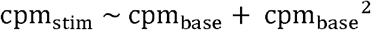

For approximately one third of genes (*n* = 466), the residuals from both of these regressions deviated significantly from the assumptions of normality and/or homoscedasticity, and a non-parametric Kendall–Theil linear regression was fitted instead. Regression fits varied from gene to gene (R^2^ ranging from <0.001 to 0.85).

#### Inter-individual variability in response

Inter-individual variability in response was quantified by comparing the coefficient of variation (CV) for baseline abundances across individuals (CV_base_) and the CV for stimulated abundances across individuals (CV_stim_). As response is defined as the difference between baseline and stimulated abundance, a large difference in their CVs, either

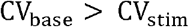

or

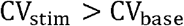

indicates a high level of variability in response. A relationship between gene-wise mean expression levels and CV is typically found in RNASeq data, with low mean transcript abundance being strongly associated with high variability^37^. As we restricted our analysis to informative genes only, excluding those genes with low mean abundance, it was not necessary to account for this relationship (Supplementary Fig. 2). Asymptotic tests for the equality of CVs were run using the cvequality package. All *p*-values were corrected for multiple testing using the Benjamini-Hochberg method^38^.

#### Functional annotation

Functional enrichment analyses were run using The Database for Annotation, Visualization and Integrated Discovery (DAVID) version 6.8^39,40^. Benjamini-Hochberg corrected *p*-values and gene counts are reported alongside ontology terms, including Kyoto Encyclopedia of Genes and Genomes (KEGG) pathways to indicate their level of enrichment^41–43^.

#### Age-specific analysis

In order to begin to investigate the relative importance of genetic variation versus prior stimulation for shaping patterns of variation in immune response, the same analysis was performed separately on juvenile and mature voles. As we had more samples from mature voles (*n* = 43) than juvenile voles (*n* = 19), we randomly sampled the mature population (with replacement) 1000 times and averaged across these samples. The number (juveniles) or mean number (matures) of genes in each of these age classes is presented in the text.

## Acknowledgements

The authors wish to thank those involved in obtaining and processing samples from the field: Rebecca Turner, Lukasz Lukomski, Stephen Price, William Foster, Ann Lowe and Anna Thomason. They also wish to thank the Forestry Commission for access to the study sites and the Centre for Genomic Research at the University of Liverpool for sequencing samples.

## Author contributions

M.B., J.E.B., J.A.J. and S.P. designed the study. E.A. undertook the stimulatory assays.

K.M.W. analysed the data. All authors wrote the manuscript.

## Competing interest statement

The authors declare no competing financial interests.

