## Supplementary information for "Decomposing variation in immune response in a wild rodent population"

**Supplementary Fig. 1 Comparison of expression levels for four genes (*Cd8a, Arg1, Foxp3 and Il1b)* deduced from RNASeq and two-step reverse transcription quantitative real time PCR (Q-PCR) performed on the same control (baseline) samples.** Counts per million per gene (RNASeq) are plotted on the x-axes, and, expression levels relative to the house-keeping gene, *Ywhaz*, (Q-PCR) are plotted on the y-axes. See Jackson *et al.*[1] for detailed methodology.

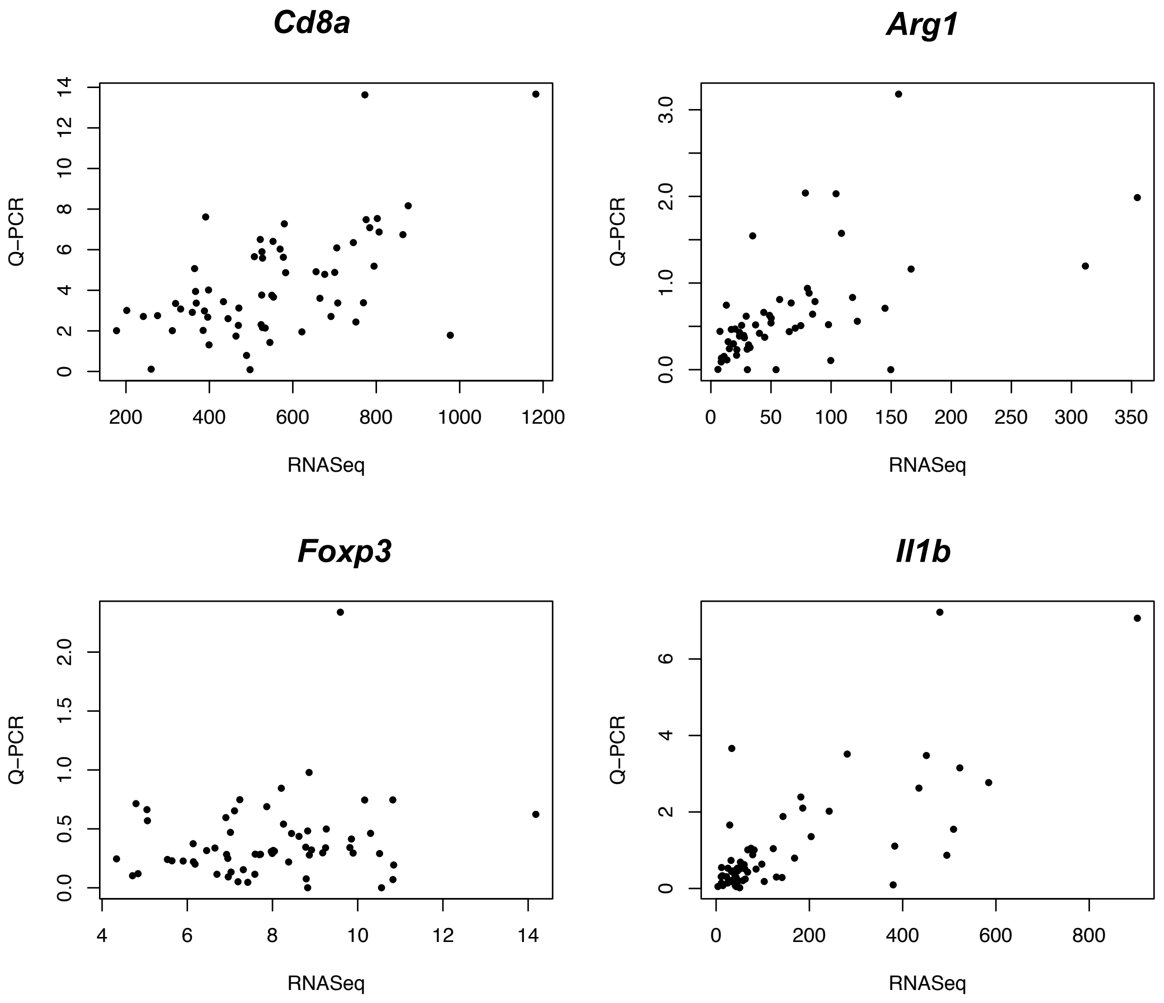

**Supplementary Fig. 2 Relationship between gene-wise means and coefficients of variation (CV) for expression levels in (a) control (baseline) samples and (b) stimulated samples.** Weakly expressed genes were omitted from the analysis. A threshold of 200 counts per million (cpm) in baseline samples and/or stimulated samples (indicated by dashed line) was chosen because this is the region in which the mean-variance relationship asymptotes i.e. a gene's variance becomes independent of its mean expression level, and other variables used to categorise genes also become independent of mean expression levels (as tested by Spearman's rank correlation tests).

**a**

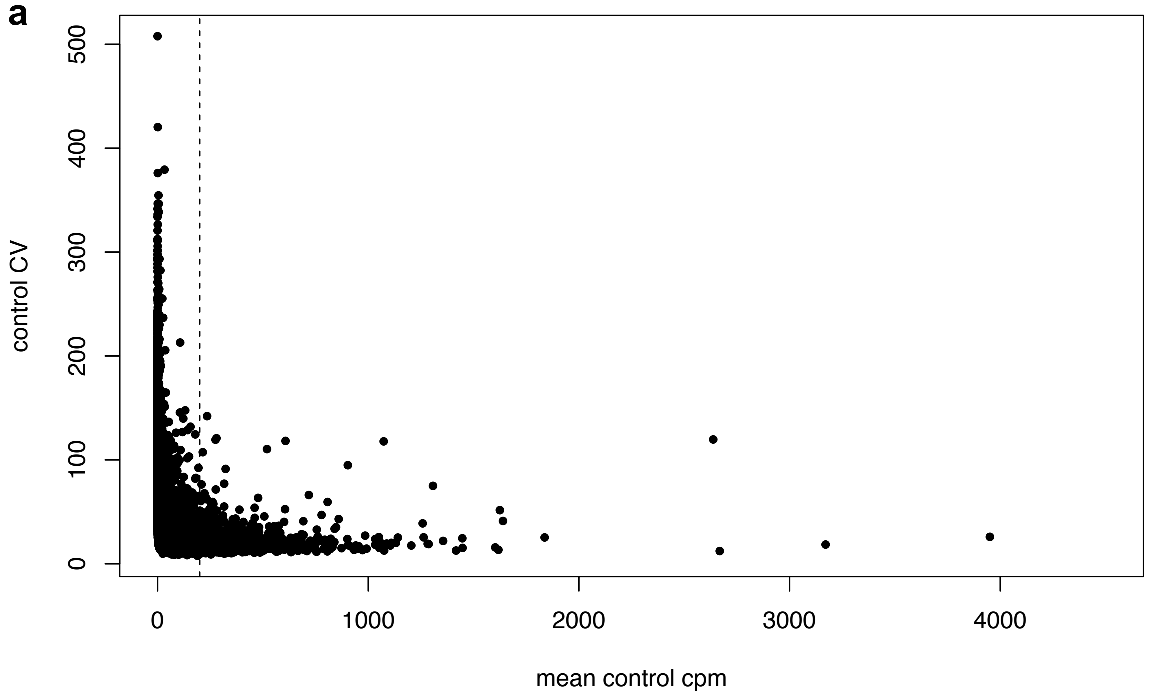

**b**

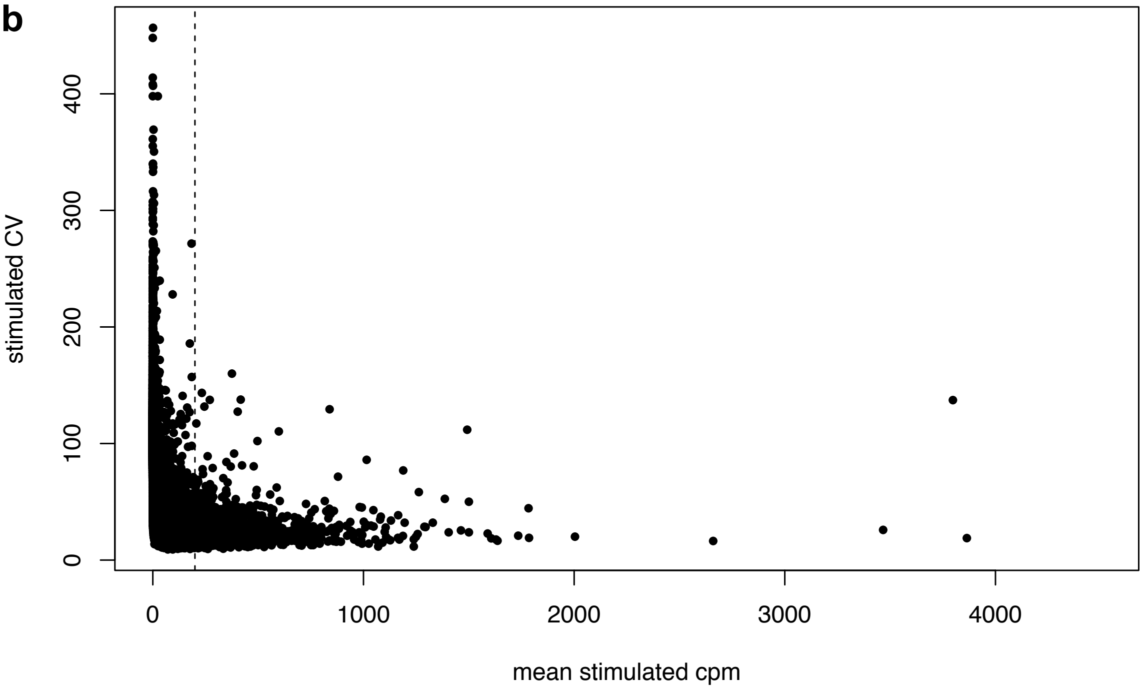
